# High-dimensional population codes reveal interpretable and diverse features underlying visual perception

**DOI:** 10.64898/2026.09.05.748439

**Authors:** Habon Issa, Sunny Liu, Jona Ballé, David Klindt

**Affiliations:** Cold Spring Harbor Laboratory, Cold Spring Harbor, NY, USA; Department of Electrical and Computer Engineering, New York University, New York, NY, USA

## Abstract

Despite large-scale recordings, neuroscience has identified representations for only a small fraction of visual features distinguishable by humans. This is increasingly attributed to mixed selectivity, where individual neurons respond to multiple stimuli, requiring population-level readouts of human-recognizable (‘interpretable’) representations. To identify interpretable representations at scale, we 1) computationally extract population codes from macaque electrophysiology datasets spanning V4 and IT cortex and vision models and 2) introduce an automated method to measure representation interpretability and diversity, then isolate a set of unique, meaningful features encoded by neurons versus populations. Across datasets, populations represent more interpretable, diverse features than neurons. This advantage grows with the number of sampled neurons and image diversity. Finally, we demonstrate through targeted ablations that interpretable representations play a causal role in downstream behavior. Overall, our framework enables a more comprehensive account of the features represented across biological and artificial visual systems and their contributions to visual perception.

## Main

A central aim for neuroscience is to understand how the neural code gives rise to perception and behavior. The tradition of vision research has been to approach this question one neuron at a time; characterizing the selectivity of single cells to hand-picked stimuli from oriented bars [1] to faces and hands [2, 3]. This approach embodies Barlow’s single-neuron doctrine [4], in which each neuron is ‘interpretable’ because it responds to a ‘trigger feature’ consistent enough that scientists can understand it. The doctrine has served as a practical discovery program for vision science: record more neurons and add each newly interpretable one to the field’s anthology to advance our understanding of the features that underlie perception. While recording technology has since delivered an exponential increase in the number of neurons we can characterize [5, 6], the anthology has not grown in step.

There is growing consensus that this is due to mixed selectivity [7–10], where neurons respond to multiple human-recognizable features in ways that resist the trigger-feature description the doctrine assumes, requiring population-level readouts. This is supported by evidence that population activity in V4 and IT [11, 12] cortex can collectively decode more recognizable features than the individual neurons are selective for. Moreover, IT population decoding is consistent with human object recognition performance [13], directly linking population codes in this region to perception. While this establishes populations as the relevant unit of analysis, decoding alone leaves the underlying neural code largely uncharacterized. To understand the atoms of perception at scale, the field needs data-driven discovery suited to uncover new feature representations in visual cortex.

In the last decade, dimensionality reduction methods were perhaps the most ubiquitous data-driven tools to extract meaningful representations from large-scale neural data [14, 15]. While they excel at extracting population codes for genuinely low-dimensional behaviors [6], this approach may be at odds with visual representations that tend to be high-dimensional [16]. Alternatively, the recently popularized sparse autoencoders (SAEs; 17, 18) decompose neural activations into sparse population codes. Their number can exceed the number of neurons, allowing SAEs to identify a wealth of interpretable representations in large language models [19–21] and other domains [22]. However, their utility for biological neurons remains underexplored. Finally, template matching [23–25] defines a series of canonical population-wide responses (‘templates’) to pre-defined stimulus classes, then computes the distance of each neural response vector to a given template to construct distance-based population codes. This has been successfully applied to visual cortex [23] and, like SAEs, template matching can identify more population codes than neurons. Whether performing template matching unsupervised is as successful is unclear.

Even with scalable representation learning, evaluation remains a bottleneck. Automated interpretability metrics are well-established for language [26, 27], but vision studies have traditionally used human ratings [22] and human psychophysics [28] to measure the visual and semantic consistency of neurons’ preferred images. We previously introduced the first automated interpretability task for vision which quantifies the separability of a neuron’s maximally and minimally exciting images [29]. However, symmetric metrics penalize locally tuned [30, 31] and sparsely activating [32] units whose neural responses to similar inputs can fall on opposite ends of the tuning curve, motivating an asymmetric alternative. Additionally, no existing automated interpretability methods quantify representation diversity operationalized as the difference between image preferences.

In this work, we bridge data-driven population extraction with automated evaluations to accelerate interpretable visual representation discovery. We introduce an asymmetric automated interpretability task suited for non-negative, sparse activations and a complementary measure of representation diversity. With these metrics, we ask whether PCA, SAEs or unsupervised template matching (UTM) learned on artificial vision networks [33, 34] and macaque electrophysiology datasets spanning IT and V4 [13, 35] yields more favorable scaling of interpretable representation discovery with dataset size than single-neuron analysis. We find that both SAE and UTM codes are more interpretable and diverse than neurons, providing the first systematic evidence that high-dimensional population analysis improves how feature discovery scales in models and brains. Finally, we demonstrate precise behavioral interventions can be achieved with this expanded anthology of interpretable representations. Collectively, our results establish a novel framework for a more comprehensive understanding of the neural code and how it shapes visual perception.

### Visual representation discovery and evaluation

We pragmatically define a visual feature as an image attribute (e.g., color, object category, texture), and a visual representation as the neural response associated with that feature. Mixed selectivity [8, 10], the tendency for single neurons to respond to multiple features, is prevalent in both visual cortex and artificial neural networks [7, 9, 36]. Fig. 1 illustrates our general approach to interpreting the neural code by extracting population codes from mixed-selective neurons: if two neurons (*Y*_1_, *Y*_2_) define axes in an activation space, their joint activation levels specify directions (population codes *Z*_1_-*Z*_3_) that are each more interpretable than either neuron alone (Fig. 1a, left). We evaluate the interpretability and diversity of each representation by examining the consistency of a representation’s preferred images (set sizes in Fig.1a, right) and the uniqueness of the representation’s preferences compared to other units (arrows between sets in Fig. 1a, right) respectively. We employ three approaches to extract population codes for subsequent interpretability evaluation. The first is principal component analysis (PCA) which finds a set of orthogonal directions to project the data onto (Fig. 1b). The second is sparse autoencoders (SAEs) (Fig.1c), which decompose neural activations into sparse latent codes. The third is an unsupervised variant of template matching (UTM; Fig.1d), which computes population activations by measuring distances between canonical population responses (K-means cluster centers or ‘templates’) and individual image responses.

**Fig. 1:**
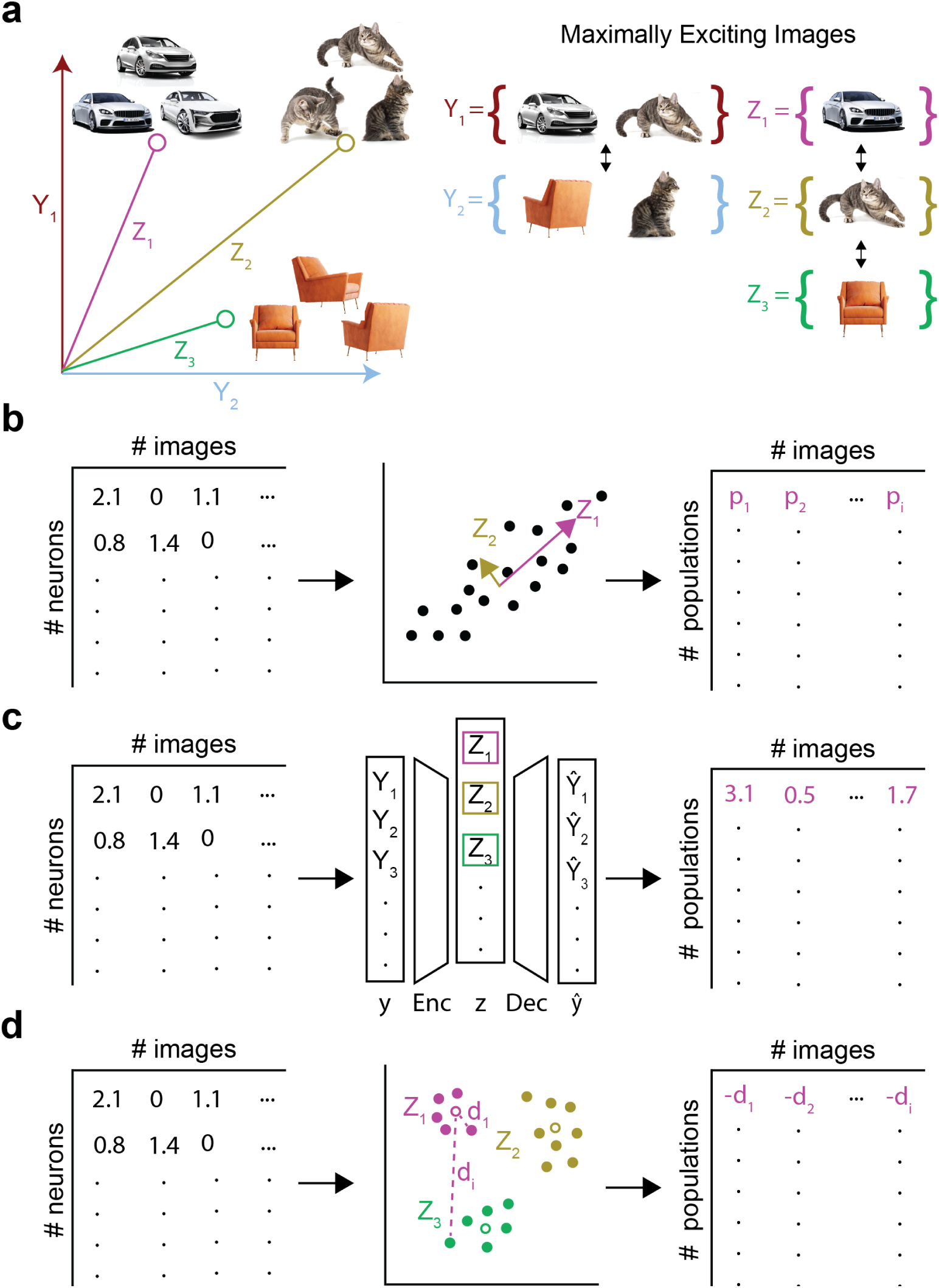
Visual representation discovery and evaluation. **a**, Left: An activation space defined by two neurons (*Y*_1_*, Y*_2_), that each respond maximally to multiple visual features. We aim to identify population codes, directions in the neural activation space (*Z*_1_ *− Z*_3_), that align with single visual features (e.g., cat). Right: Sets which define the number of distinct visual features among each representation’s top 3 maximally exciting images (MEIs). We operationally define interpretability as the consistency of a representation’s MEIs (set sizes), and diversity as the separability of two representation’s preferred features (double headed arrows). **b-d,** Population code extraction methods. We transform single- and multi-unit responses to images into population responses using three independent methods: principal component analysis (PCA), sparse autoencoders (SAEs) and unsupervised template matching (UTM). **b,** PCA. After identifying Z orthogonal directions in the data (arrow 1), each row of the population response matrix is populated with the projections of each column of the original dataset onto a single principal component (arrow 2). **c,** SAEs. We identify Z sparse (L1), reconstructive (MSE) codes from the original data (arrow 1), filling each row of the population response matrix with a single latent’s image responses (arrow 2). **d,** UTM. With K-means, Z cluster centers (open circles) represent canonical population-wide responses to images (arrow 1; filled circles represent columns of the original data). Each row of the population response matrix is comprised of the negative Euclidean distances between one cluster center and each original datapoint (arrow 2).

For the main text, we perform this analysis on representative subjects from two public primate electrophysiology datasets, specifically: IT cortex responses to faces and objects from Vinken et al. [35] and IT and V4 responses to grayscale objects and faces under various identity-preserving transformations from Majaj et al. [13]. We additionally analyzed the responses of three computer vision models, ResNet-50 (layers 3 and 4) [34], CLIP ViT-B/16 [33], and DINOv2 [37] to 10,000 CIFAR-100 images [38]. DINOv2 responses to Vinken et al. [35] images are included in a subset of experiments.

In the extended data section, we report additional subjects’ results (159 IT units, N=3 macaques from [35]; 110 IT units, N=1 macaque from [13]) provided a minimum of 30 units were recorded from each subject. Finally, we report results from AM neuron responses to grayscale face images (159 neurons, 2100 images, N=2 macaques from [39]).

### Populations represent more interpretable visual features than neurons

We evaluate interpretability with an Odd-One-Out task performed *in silico* (Fig. 2). This was first proposed for interpretability measurement as a word intrusion task, where the odd-word-out can easily be detected among an otherwise semantically consistent list [40]. For our task, we define a per-representation threshold by computing the average pairwise similarity of each neuron or population’s MEIs using Dream-Sim, a model trained on human perceptual judgments [41]. All remaining images are intruders, and a point is scored each time an intruder’s average pairwise similarity to the MEIs falls below the task threshold (Fig. 2a). Our previous symmetric version of this task [29] was found to have high correspondence with human psychophysics judgments, though we opted for an asymmetric design to accommodate sparsely activating units (e.g., SAE latents). Fig. 2c shows relatively high and low scoring neurons in IT cortex from the Vinken et al. [35] dataset, demonstrating that our task score is a valid proxy for interpretability operationalized as consistency among a unit’s preferred images.

**Fig. 2:**
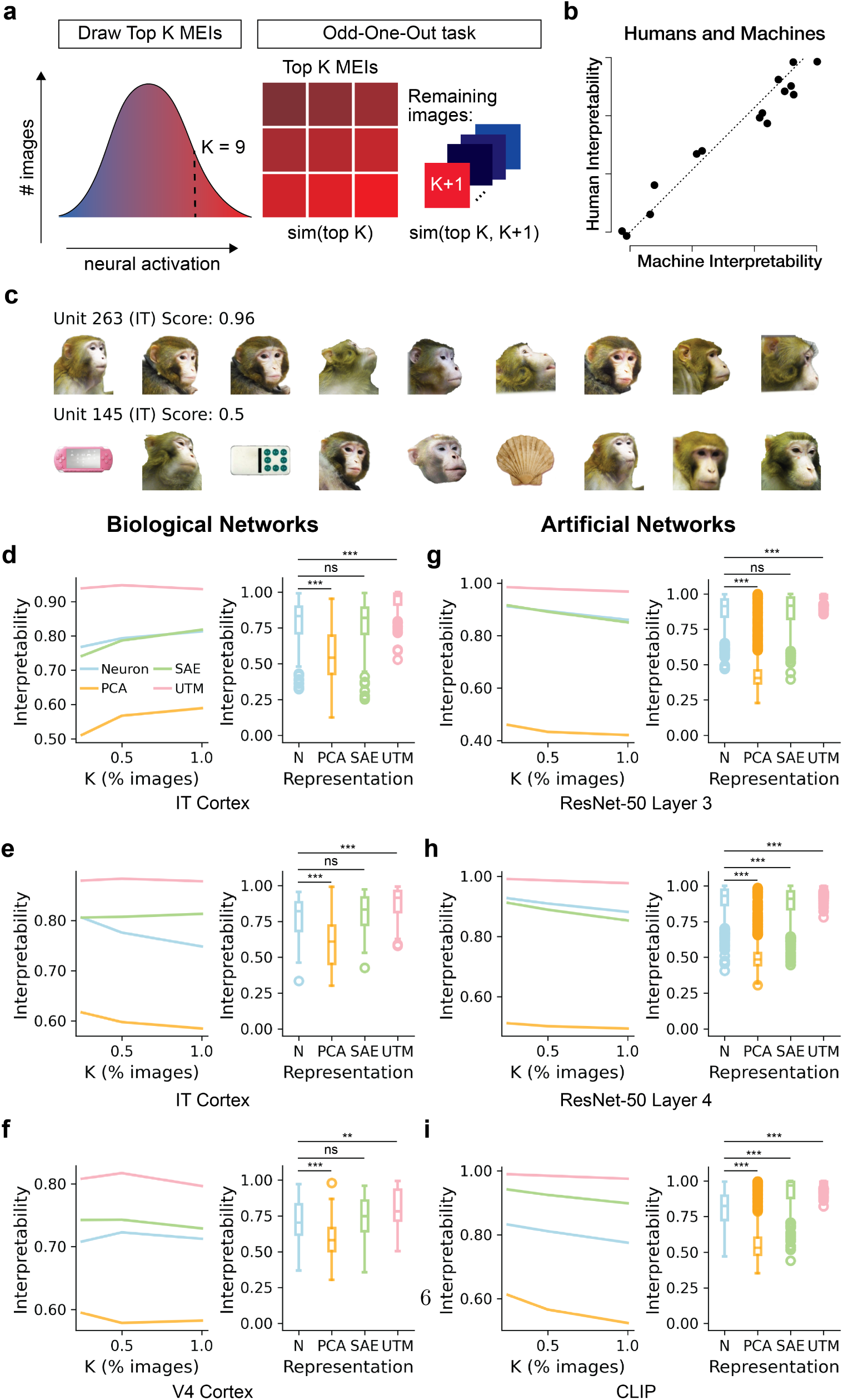
Interpretability metric and results. **a**, The Odd-One-Out task measures per-unit (neuron or population) interpretability. Given K MEIs (left), we compute the fraction of trials where non-MEI images are less similar to the MEIs than the MEIs are to each other (right), using DreamSim [41] as the image similarity metric. **b,** High correspondence between a symmetric version of the Odd-One-Out task and human psychophysics judgments on the same task. Reproduced with permission from Zimmermann et al. [42]. **c,** 9 MEIs and corresponding Odd-One-Out scores for interpretable (top) and uninterpretable (bottom) IT neurons from the Vinken et al. [35] dataset. **d-i,** Interpretability scores. Left plots show average interpretability scores at each value of K (as a percentage of all images in the dataset). Right plots additionally average across all values of K. **d-f,** Visual cortex results. PCA, SAE, and UTM codes were less, equally and more interpretable than neurons, respectively. Data: Vinken et al. [35] (**d**) Majaj et al. [13] (**e,f**). **g-i,** Vision model results. PCA codes were less interpretable than neurons. SAE codes were equally as interpretable as neurons for ResNet-50 layer 3 (**g**), less interpretable than neurons for ResNet-50 layer 4 (**h**), and more interpretable than neurons for CLIP (**i**). UTM codes were consistently more interpretable than neurons. ns = Not Significant, ** = *P <* 0.01, *** = *P <* 0.001, Kruskal-Wallis with Dunn’s post hoc test. Significance markers indicate comparisons to the neuron baseline; all comparisons are reported in Extended Data Tables 1,2.

On average, PCA codes are less interpretable than neurons across biological and artificial neural networks (Figs. 2d-i). SAE codes had equivalent performance to the biological and ResNet-50 layer 3 neurons (Figs. 2d-g), underperformed ResNet-50 layer 4 neurons (Fig. 2h), and outperformed CLIP neurons (Fig. 2i). UTM codes are more interpretable than neurons across datasets (Figs. 2d-i). UTM also improved interpretability relative to SAEs on all datasets except the V4 recordings (Extended Data Tables 1,2). These findings suggest that the most interpretable visual representations are not aligned with single neurons or orthogonal directions of highest variance in the data (i.e., PCA).

Receptive fields may confound our single-neuron results. To test their contributions, we performed spatially restricted Odd-One-Out measurements, finding no difference in interpretability scores for V4 neurons (Extended Data Fig. 1a). V4 interpretability did increase when measuring Odd-One-Out with the Wasserstein Distortion [43], a metric which measures local image similarity using VGG-19 feature maps (Extended Data Fig. 1b). In contrast, spatially restricted DreamSim measurements increased the interpretability of AM neurons known to respond to local but readily recognizable concepts (Extended Data Fig. 1e). This supports the notion that V4 neurons are genuinely more ‘entangled’ [13] than the representations typically favored by DreamSim.

Together, our results demonstrate that high-dimensional population representations are more interpretable than neurons for both artificial and biological networks. This effect is not merely due to differences in receptive field sizes, and is strongest with UTM. Note that the relative level of interpretability may depend on the image dataset. Thus, it is comparable between the IT and V4 regions of the Majaj et al. [13] dataset (Fig. 2e, f), and between artificial vision network responses to CIFAR-100 (Fig. 2g-i), but not across datasets.

### Populations represent more diverse visual features than neurons

The Odd-One-Out task does not distinguish between interpretable representations that are redundant (e.g., face cells coding identical category information) and unique (e.g., ResNet-50 layer 4 neurons mapping to distinct object categories). We therefore introduced a second metric, feature diversity, which captures the redundancy of feature preferences across units (neurons or populations). This Cross Odd-One-Out diversity task measures how unique a reference unit’s MEIs are compared to all other units. For each pairwise comparison, we calculate how often an intruder image (from another unit’s MEIs) is correctly identified as not belonging to the reference unit’s MEIs (Fig. 3a) based on DreamSim embeddings. High scores indicate distinct feature preferences between units.

**Fig. 3:**
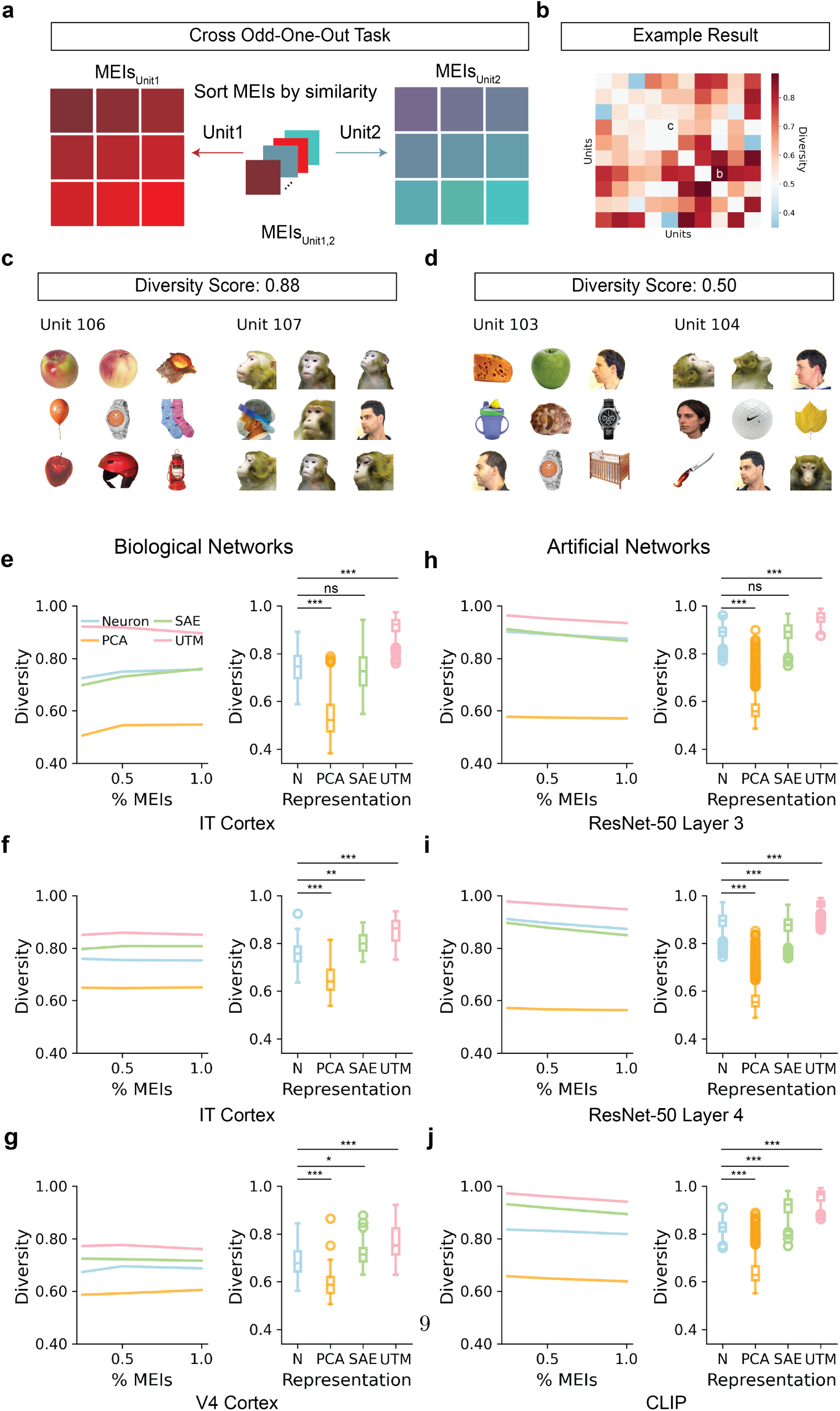
Diversity metric and results. **a**, The Cross Odd-One-Out task measures the average stimulus preference differences across all pairwise unit comparisons (neurons or populations). Given K MEIs from each of two units, we compute the fraction of trials where an MEI is more similar to the other MEIs of its own unit than to the MEIs of the opposing unit, using DreamSim as the image similarity metric. **b,** Pair-wise comparisons among 10 example units. 0.5 represents chance level. **c,d,** 9 MEIs and Cross Odd-One-Out scores for a pair of units with distinguishable (**c**) and indistinguishable (**d**) feature preferences. Images from Vinken et al. [35]. **e-j,** Diversity scores. Left plots show diversity scores averaged across N units at each value of K (as a percentage of all images in the dataset). Right plots show the same data, averaging Cross Odd-One-Out scores across values of K. **e-g,** Visual cortex results. PCA codes are less diverse than neurons. SAE codes are equally diverse for Vinken et al. [35] IT neurons (**e**) and more diverse for IT and V4 neurons of the Majaj et al. [13] dataset (**f,g**). UTM codes are more diverse than neurons. **h-j,** Vision model results. PCA codes are less diverse than neurons. SAE codes are equally as diverse as ResNet50 layer 3 neurons (**h**), less diverse than ResNet-50 layer 4 neurons (**i**), and more diverse than CLIP neurons (**j**). UTM codes are more diverse than neurons. ns = Not Significant, *** = *P <* 0.001, Kruskal-Wallis with Dunn’s post hoc test. Significance markers indicate comparisons to the neuron baseline; full pairwise comparisons are reported in Extended Data Tables 3,4.

The Cross Odd-One-Out results mirror our Odd-One-Out findings, but with generally larger effect sizes (Extended Data Table 4). PCA codes are less diverse than neurons across datasets (Fig. 3e-j). For biological neurons, SAE codes have equivalent performance to the Vinken et al. [35] IT neurons (Fig. 3e) and outperform the IT and V4 neurons of the Majaj et al. [13] dataset (Figs. 3f, g). SAEs have equivalent performance to the ResNet-50 layer 3 neurons (Fig. 3h), under-perform the ResNet-50 layer 4 neurons (Fig. 3i), and are more diverse than the CLIP neurons (Fig. 3j). UTM representations are consistently the most diverse compared to both neurons and SAE codes (Extended Data Table 4).

Given the consistent performance of UTM codes on both representation evaluation tasks, subsequent analyses compare neurons to UTM codes directly. To confirm that our UTM findings reflect population synergy, we report control results for the Odd-One-Out and Cross Odd-One-Out tasks in Extended Data Fig. 2. Our aim is to remove synergistic information from the population without reducing baseline interpretability to chance levels. To accomplish this, we construct a null IT population (N=85 neurons) by selecting the most interpretable neuron, duplicating it 85 times, and adding independent Gaussian noise to each copy, with noise magnitude matched to the standard deviation of the original 85 IT neurons. UTM applied to this null population decreases interpretability (Extended Data Fig. 2c) and increases diversity (Extended Data Fig. 2d) without matching the diversity of the original data. Thus, our UTM results point to genuine synergy in the neural populations being examined. Collectively, these results demonstrate that high-dimensional population representations are more diverse than neurons, indicate that diversity is a primary advantage of population coding in later stages of visual cortex, and highlight that population synergy is required for neural systems to represent features not encoded at the single-neuron level.

### Accelerating interpretable representation discovery

Individually, the Odd-One-Out and Cross Odd-One-Out tasks highlight that on average, UTM populations encode more interpretable, diverse visual features than neurons. Using a greedy algorithm on the combined task scores, we can report the precise number of distinct visual features discovered by UTM versus neurons (Fig. 4a).

**Fig. 4:**
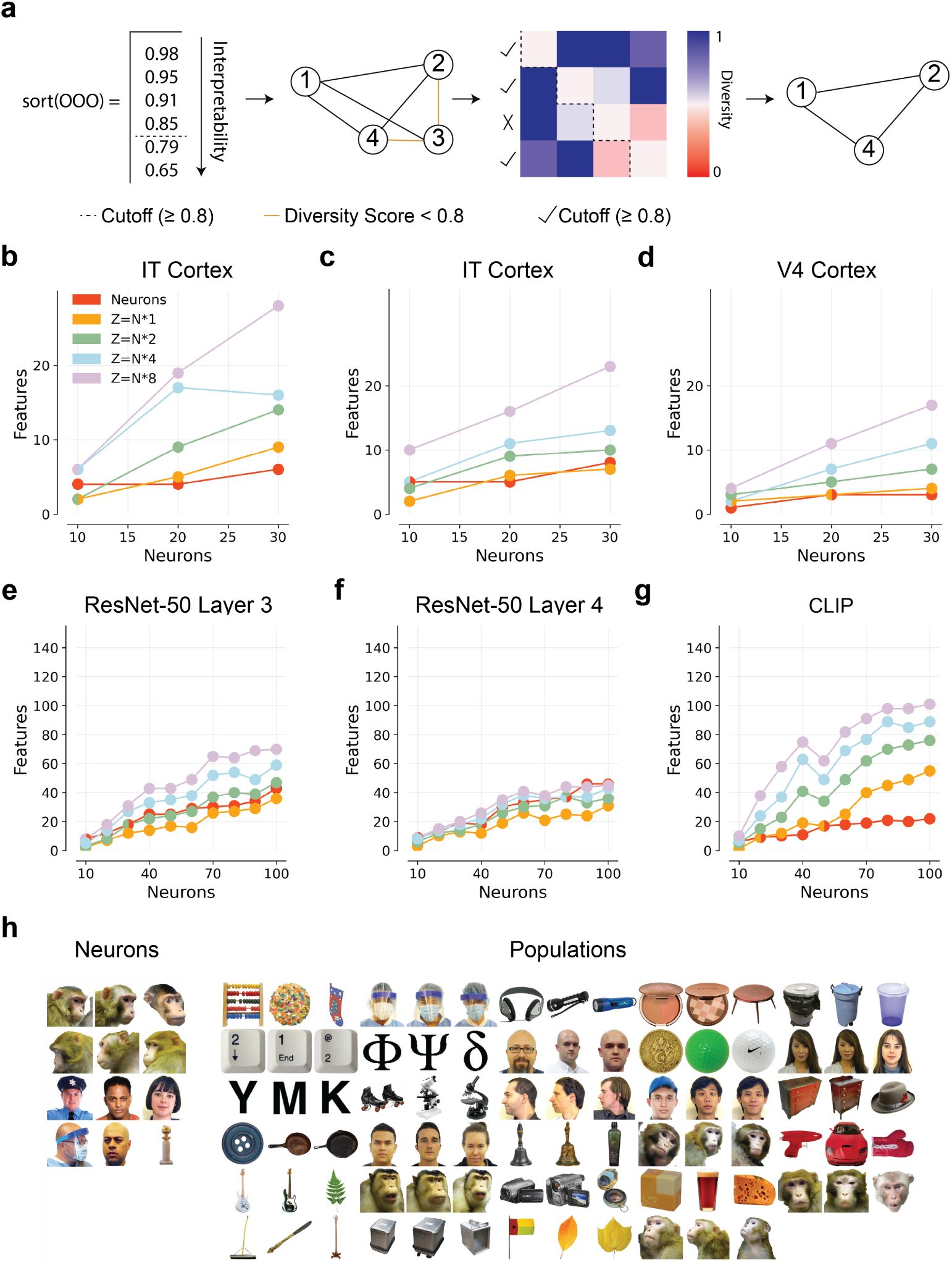
Scaling laws of visual feature discovery. **a**, Algorithm used to define a unique set of interpretable features. We sort all units in order of interpretability, retaining only those that have Odd-One-Out scores *≥* 0.8 (arrow 1). We append these units to a growing list if their Cross-Odd-One-Out score with any unit on the list is *≥* 0.8 (arrow 2). This leaves us with approximately the most interpretable, diverse set of possible units (arrow 3). We plot the list size (number of discovered features) in subsequent subpanels. **b-d,** Plots showing the number of unique, interpretable representations discovered per biological dataset. For each dataset, we compute this number for a sample of N=10, 20 and 30 neurons (red points). For each number of neurons, we perform UTM at expansions of 1*N*, 2*N*, 4*N* and 8*N*, computing the number of unique interpretable representations for each expansion factor (orange, green, blue and purple points respectively). Data from Vinken et al. [35] dataset (**b**), Majaj et al. (IT) [13] dataset (**c**) and Majaj et al. (V4) dataset [13] (**d**). **e-g,** Plots showing the number of unique, interpretable representations discovered per model. For each dataset, we compute this number for a sample of N=10-100 neurons in intervals of 10 (red points). For each number of neurons, we perform UTM at expansions of 1N, 2N, 4N and 8N, computing the number of unique interpretable representations for each expansion factor (orange, green, blue and purple points respectively). Data from ResNet-50 layer 3 (**e**), ResNet-50 layer 4 (**f**) and CLIP (**g**). **h,** 3 MEIs for four features discovered with neurons (left) versus 29 features discovered with UTM codes (right). Both neuron and UTM results are from N=30 subsampled IT neurons (**b**) and Z = 8N. Images from Vinken et al. [35].

Starting with *N* neurons, we sort units in descending order of interpretability, and implement a cutoff for all units with an Odd-One-Out score below 0.8. The remaining units are interpretable but potentially redundant. We next append each unit to our final list only if its pairwise Cross Odd-One-Out score with every existing list member is *≥* 0.8. This ensures we retain a list of the most interpretable instances of each visual feature. We report the length of this list in Fig. 4 as the number of discovered interpretable features.

To probe the effect of neural dataset size on feature discovery, we subsample neurons *N* in intervals of 10 (10 *−* 30 for biological networks and 10 *−* 100 for artificial networks). For each value of *N*, we perform UTM with a variable number of clusters 1, 2, 4, and 8 times *N* before applying the greedy algorithm described above. The maximum values of *N* for the biological and artificial networks are restricted such that the number of images *I* in a given dataset exceeds 10 times the maximum number of UTM codes *Z* (*I >* 10*Z*) There is one exception noted below for Vinken et al. [35], where *I ≈* 6*Z* when *Z* = 8*N*.

On all but one dataset (ResNet-50 layer 4), UTM yielded more distinct interpretable features than neurons. This effect grew with the number of UTM codes *Z* (Fig. 4b-g). ResNet-50 layer 4 is likely the exception since neurons in this layer are highly category-aligned, limiting their potential for cross-category synergy. This is consistent with the relatively low level of discovered features in layer 4 (Fig. 4f) compared to layer 3 (Fig. 4e) and CLIP (Fig. 4g), which both show evidence of superposition by representing more features than neurons [36]. While biological and artificial neural network results are not directly comparable, we next asked if the gap in image diversity between datasets may contribute to apparent differences in synergy.

Indeed, while increasing the number of images (but not categories) did not increase the number of recovered feature representations (Extended Data Fig. 3a), increasing the number of image categories did (Extended Data Fig. 3b).

In Fig. 4h we show the full set of features recovered from neurons vs UTM codes at *Z* = 8*N* from one Vinken et al. [35] subject. While neurons consistently appear to encode human and monkey face categories (Fig. 4h, left), we discovered additional UTM codes for concepts resembling objects, identity, and personal protective equipment (PPE; Fig. 4h, right). In this example, the discoveries made with populations outnumber neurons 29:4. Given the size of the feature anthology for vision models, and finding that most biological codes are category-aligned, we graphically report the vision model anthology as the number of category-aligned concepts recovered from the models in Extended Data Fig. 4. Specifically, we quantify the number of UTM or neuron codes passed through the greedy algorithm that contain majority MEIs (7/10) corresponding to a single coarse (Extended Data Fig. 4, top) or fine (Extended Data Fig. 4, bottom) label from the CIFAR-100 dataset. This analysis is performed at each number of sampled neurons, and the number of recovered CIFAR-100 labels is measured cumulatively. By *N* = 100, all coarse category labels are recovered from each vision model using UTM codes. CLIP UTM codes also recover nearly every fine CIFAR-100 category label by *N* = 100, while UTM codes of ResNet-50 layers 3 and 4 yield roughly half of the labels. For both coarse and fine labels, the gap between ResNet-50 layer 4 and UTM codes is minimal, indicating pre-existing category selectivity, rather than population synergy, is driving label recovery in this layer.

These results demonstrate that our computational framework achieves more favorable scaling between the number of recorded neurons and the biological insights they provide. There is no apparent limit to this scaling, indicating that larger image and/or neural datasets will increase the gap between discoveries made from neurons compared to populations. Finally, most discovered representations appear to encode mid- to high-level concepts including identity and object category. We attribute this to the combination of object-centric images with DreamSim, and do not rule out the possibility that interpretable lower level or domain-general feature representations are encoded by UTM populations.

### UTM codes enable precise behavioral control

One goal of interpretable representation discovery is to understand what features are important for perception and behavior. To address whether features discovered using our framework provide these insights on downstream tasks, we start by measuring model classification performance before and after single neuron or UTM population ablations (Fig. 5). We perform this analysis with DINOv2, a vision transformer with patch-based activation maps [37]. This affords us post-hoc explanations for the precise feature within the MEIs that is being manipulated, for example the outer portion of the ‘sunflower’ feature (Fig. 5a). Fig. 5b shows a classification task among all 5 members of the CIFAR-100 ‘flowers’ superclass. Ablating the ‘sunflower’ UTM code has the strongest effect on sunflower classification, which are more frequently confused with roses and poppies. Repeating this experiment with all remaining category-aligned UTM codes discovered from DINOv2 responses to CIFAR-100 reveals that the class most associated with the ablated feature tends to be selectively affected by UTM ablations (Fig. 5c, d). In contrast, ablating interpretable neuron contributions did not change overall classification performance (Fig. 5g, h) and indicates that single neurons have a negligible effect on behavior.

**Fig. 5:**
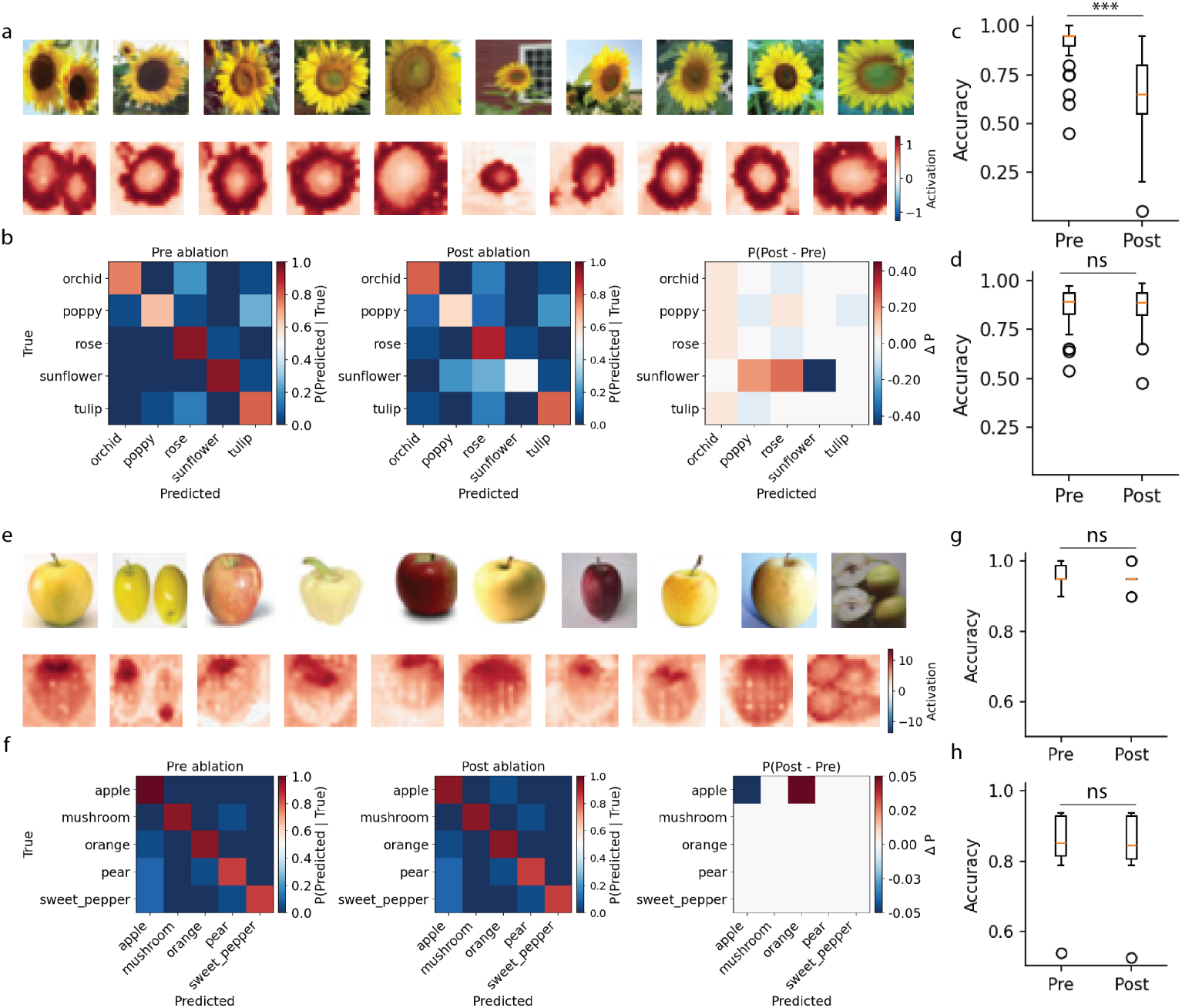
Behavioral control with DINOv2 populations. **a**, MEIs (top) and strongly activated patches (bottom) for a UTM population appearing to encode the outer portion of sunflowers. Images from Krizhevsky et al. [38]. **b,** Confusion matrix before (left) and after (middle) ‘sunflower’ feature ablation. The difference between confusion matrices (after - before) is shown on the right. **c,** Average classification error rate for the relevant class (e.g., sunflower when the ‘sunflower’ feature is ablated) pre versus post UTM feature ablation. **d,** Average classification error rate for all other classes (e.g., orchid, poppy, rose, and tulip when the ‘sunflower’ feature is ablated) pre versus post UTM feature ablation. **e-h,** Same as **a-d** but with neurons instead of UTM codes. Images from Krizhevsky et al. [38].

To perform an analogous experiment on the biological data, we 1) introduce more precise receptive field estimation to generate post-hoc ablation explanations and 2) train a linear classifier to test ablation effects on the static visual cortex responses. We perform receptive field estimation by using regression to predict DINOv2 UTM code responses from all (Fig. 6a-d) or one (Fig. 6e-h) of the Vinken et al. [35] neurons. Linear combinations of biological neurons with *R*^2^ *≥* 0.6 serve as our biological population codes, while individual neurons with *R*^2^ *≥* 0.6 are retained for the single neuron analysis. We subsequently test the effects of population or single neuron ablations on the performance of a linear classifier trained to predict Vinken et al. [35] labels from the original IT neurons. Population ablations decrease classification accuracy for the ablation-relevant class (Fig. 6c) and produce relatively small increases in classification accuracy for other classes (Fig. 6d). Neuron ablations did not change classification accuracy for the ablation-relevant class (Fig. 6g) and produced a small increase in classification accuracy for other classes (Fig. 6h).

**Fig. 6:**
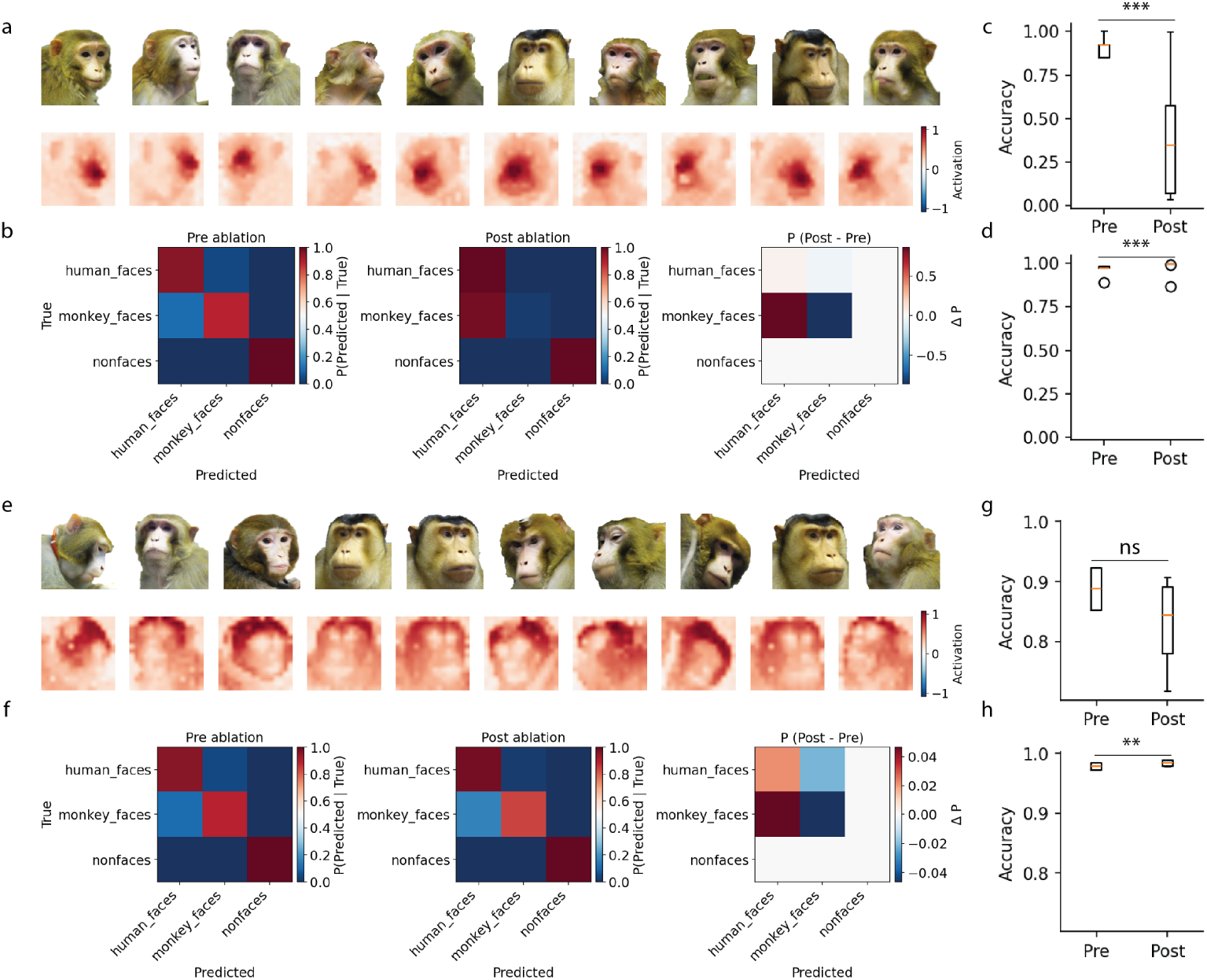
Behavioral control with IT cortex populations. **a**, MEIs (top) and strongly activated patches (bottom) for a UTM population appearing to encode macaque noses. Images from Vinken et al. [35]. **b,** Confusion matrix before (left) and after (middle) the ‘macaque’ feature ablation. The difference between confusion matrices (after - before) is shown on the right. **c,** Average classification error rate for the relevant class (e.g., macaques when the ‘macaque’ UTM code is ablated) pre versus post UTM feature ablation. **d,** Average classification error rate for other classes (humans and nonfaces) pre versus post UTM feature ablation. **e-h,** Same as **a-d** but with neurons instead of UTM codes. Images from Vinken et al. [35].

Overall, these results demonstrate that UTM codes have the most precise, interpretable, and potent effects on downstream behavior. It’s worth noting that brain- to-model prediction presents a significant bottleneck for the IT cortex analysis (Fig. 6), particularly for single neurons. Nonetheless, we find ‘reverse predictive’ approaches (brain-to-model mappings; see [44]) to receptive field localization a promising future direction given the general convergence between model and brain representations [45, 46]; and anticipate that the yield of this approach will increase with dataset size and diversity.

## Discussion

Since the inception of the single-neuron doctrine, Barlow [4] argued that ‘semantically coherent’ (interpretable) and ‘redundancy-reducing’ (diverse) representations together give rise to perception. There is mounting evidence that such representations exist at the population level [47], however, conventional analysis methods tend to recover far fewer interpretable features than the number of neurons recorded.

To address this, we recover high-dimensional population codes, introduce automated metrics for representation interpretability and diversity, and quantify the degree to which high-dimensional population codes store visual information obscured at the level of single neurons. We demonstrate that high-dimensional population codes represent more interpretable, diverse visual features than neurons (Figs. 2,3) and close the gap between dataset size and discovery (Fig. 4). The discovered features provide explanations for perception by enabling precise, predictable control over behavior (Figs. 5, 6). Our results demonstrate that high-dimensional representations are the relevant unit of analysis for a more comprehensive understanding visual perception, and we offer an approach to interrogate them in a way that scales with modern neural data collection.

Measuring interpretability and diversity with DreamSim evaluations of maximally exciting images (MEIs) comes with several limitations. One is that MEIs are always assigned. For example, a face-selective unit’s MEIs may be ‘apples’ in a dataset without faces. Future work in this direction may aim to have larger, or more efficiently sampled, image datasets. Another is that MEIs themselves are not features, so images with sufficient irrelevant or inhibitory [48] content may fall outside of a unit’s MEIs despite containing a trigger feature for that unit. Thus, natural extensions of our pipeline include incorporating receptive field estimation (Extended Data Fig. 1), object segmentation [49], attribution maps [50], per-patch activations (Figs. 5,6), or exploring alternative image similarity metrics (Extended Data Fig. 1).

Limitations of MEI-based analysis with DreamSim may also contribute to the difference between SAE and UTM performance. Qualitatively, SAE latents appear to encode more atomic, compositional features. Therefore, extending our pipeline may eventually reveal SAEs exceed the performance of UTM codes, so we withhold any conclusions about which of the two codes is better overall. What is clear is that both codes outperform neurons and PCA (Figs. 2,3), the extent of which supports superposition and mixed-selectivity beyond a simple axis rotation (Fig. 4). Critically, superposition (representing more features than neurons) requires representations to be non-orthogonal [36], a condition that PCA is not designed to capture. While the existence of superposition in biological networks remains unproven, plenty of pressures to make mixed-selective representations non-orthogonal exist, including generalization via continuous manifold structure.

Finding interpretable population representations with UTM enabled greater insights into perception. Ablating UTM features discovered using our pipeline yielded specific and interpretable changes to classification at inference time with DINOv2 (Fig. 5c,d). The same ablation strategy applied to interpretable neurons did not (Fig. 5g,h), consistent with previous findings that single neurons do not predict perceptual experience [51]. This is likely because neural systems are robust, and where single neurons represent one of many features aligned to a given concept, populations carry a more complete representation. Our population ablation results extend to the biological data (Fig. 6), although we are limited to building classifiers on the static datasets. Coupling our pipeline with methods such as two-photon holographic optogenetics [52] can answer whether our population steering methods are as effective in awake, behaving animals.

Collectively, our results show that we can obtain far more information about neural representations and their contributions to perception than previously realized. Data continues to be a bottleneck, but where previous calls for data focused more exclusively on neuron count, our results suggest that at least equal effort should be directed towards increasing stimulus diversity in way of several recent datasets [53–55]. Given the stability of populations, perhaps future studies can also reduce the number of image repetitions in favor of this diversity. Thus, we believe our contributions have not only begun to close the scale gap in neural feature discovery, but have also provided both the analytical framework and evidence that richer image datasets will yield correspondingly richer insights.

## Methods

### Data

#### Brain Data

Biological data is sourced from publicly available single- and multi-unit recordings from three primate electrophysiology studies [13, 35, 39]. The Vinken et al. [35] dataset consists of 449 neural responses from central inferotemporal (IT) cortex (84 single neurons and 365 multi-unit responses in and around ML and MF face-responsive patches) to 1,379 images (447 faces and 932 non-face objects on white backgrounds) from N=6 macaques. The Majaj et al. [13] dataset consists of 256 neurons (168 IT neurons, 88 area V4 neurons) responding to 3,200 grayscale images of objects and faces with identity-preserving transformations on varying backgrounds from N=2 macaques. The Chang et al. [39] dataset consists of 159 AM face patch neurons responding to 2,100 grayscale face images on plain backgrounds from N=2 macaques. All neural responses were either pre-averaged or averaged across all available image repeats (*µ* = 9.7*, σ* = 3.4 repeats for Vinken et al. [35]; 28 *−* 50 repeats for Majaj et al. [13]; 3 *–* 5 repeats for Chang et al. [39]). Brain activations are subsequently normalized (mean subtracted, divided by standard deviation) and stored in N×I (number of neurons, number of images) matrices of size (449, 1379; Vinken et al. [35]), (168, 3200; Majaj et al. [13] IT) and (88, 3200; Majaj et al. [13] V4), and (159, 2100; Chang et al. [39]). For both the Vinken et al. [35] and Majaj et al. [13] datasets, only one representative subject per study per region is reported in the main text. Results for remaining subjects are reported in the Extended Data section provided that *N ≥* 30 units were recorded from them. For Chang et al. [39] results, all 159 neurons were analyzed simultaneously.

#### Model Data

We obtain activations from each of the following model layers in response to 10,000 images from batch 1 of the CIFAR-100 dataset [38]: layers 3-4 of ResNet-50 ([34]) pretrained on ImageNet1k [56] with ImageNet1k V2 weights from the PyTorch implementation [57], vision embeddings of CLIP ViT-B/16 [33], and vision embeddings of DINOv2 ViT-S/14 [37]. Figs. 5,6 identify UTM codes for DINOv2 activations on the flattened per-patch activations. Otherwise, model activations are spatially averaged to produce N x I activation matrices for further analysis.

### Population code extraction

Three independent methods transform the original NxI neural data into ZxI (number of populations, number of images) population responses to the same images: unsupervised template matching, sparse autoencoders, and principal component analysis. N=Z across all experiments except those in Fig. 4, which explicitly test whether interpretable representation discovery rates increase as a function of Z. Training procedures for each population extraction method are described below.

### Unsupervised template matching

To transform neural activations into population activations, the original N*×*I neural response matrix is clustered over columns (images) into Z clusters with K-means. In the resulting N-dimensional space, each of the Z cluster centers represents a canonical N-dimensional population-wide response to a specific visual feature (e.g., cats). The negative Euclidean distance between a single cluster center and each of I population-wide image responses fills a single row of a new Z*×*I population activation matrix.

We use standard K-means, represented by:

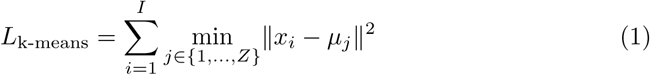

where *x_i_*represents the neural response to image *i*, and *µ_j_*represents the *j*-th of *Z* cluster centers.

### Sparse autoencoders

Sparse autoencoders (SAEs) decompose the N*×*I neural response matrix into sparse latent representations (Z*×*I). To train an SAE, the N-dimensional neural response *x* is passed into a linear encoder with a ReLU activation function, represented by:

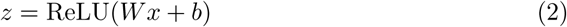

where *z* represents the SAE latents, *x* represents the input neural response, and *W* (weight matrix) and *b* (bias) are parameters learned using a combined loss function consisting of sparsity loss and reconstruction loss. The sparsity loss function, given by:

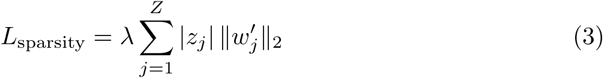

encourages the SAE to find sparse representations by penalizing nonzero SAE latent activations, weighted by each latent’s decoder column norm ‖w^’^_j_‖_2_. The strength of this penalty is controlled by the hyperparameter *λ*. The SAE learns which activations to suppress by jointly minimizing the reconstruction loss, which measures the difference between the input neural responses *x* and the reconstructed responses *x*^. The reconstruction loss is given by:

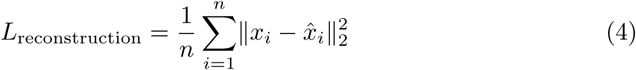

where ‖•‖^2^_2_ denotes the mean squared error across the *N* neurons, consistent with standard MSE loss implementations. The SAE decoder reconstructs input neural activity from SAE latents linearly:

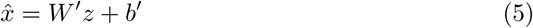

*W* ^′^ (weight matrix, with columns *w*^′^) and *b*^′^ (bias) are learned decoder parameters optimized by the reconstruction loss (Equation 4). 5-10 SAE models are trained per dataset, varying the hyperparameter *λ* between 1*e −* 2 and 1*e*1 in 5-10 logarithmically spaced steps, and using a 1x expansion factor (N=Z). The best model is selected based on a rate-distortion curve minimizing the joint MSE-L1 loss.

### Principal component analysis

Principal component analysis (PCA) finds a series of orthogonal directions of maximal variance in the original I*×*N data by first computing its covariance matrix **C**:

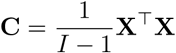

where **X** is the centered input data, and *I* is the number of images. Next, the eigenvectors and eigenvalues of this matrix are identified:

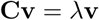

Finally, in rank order, the projection of each eigenvector onto all I population-wide responses from the original N*×*I data constitutes one row of the Z*×*I population response matrix.

### Quantifying representation interpretability and diversity

Representation interpretability and diversity are quantified with two tasks, Odd-One-Out and Cross Odd-One-Out respectively. Both tasks automate representation evaluations using one of two image similarity metrics: DreamSim [41] or the Wasserstein Distortion [43]. Image similarity metrics consider the entire image unless otherwise noted (i.e., local similarity measurements in Extended Data Fig. 1). Tasks, metrics, and local image similarity measurement procedures are described in the following sections.

### Image similarity metric: DreamSim

Trained on human psychophysics judgments, DreamSim [41] allows each representation evaluation task to be automated but human-perception aligned. The full DreamSim training procedure is described by Fu et al. [41]. Briefly, human participants were shown image triplets consisting of one reference image (Ref) and two of its distortions (A and B). Triplets for which human raters unanimously agreed on which distortion was more similar to Ref were retained for model training. During training, distances between Ref and A (d0) and between Ref and B (d1) were computed from the embeddings of an ensemble of existing state-of-the-art vision models. If humans rated distortion A as more similar to Ref, d0 was minimized in the model’s embedding space. Conversely, if humans rated distortion B as more similar to Ref, d1 was minimized in the model’s embedding space.

Before performing the Odd-One-Out and Cross Odd-One-Out tasks for the main text and Extended Data Figs. 1a (left), 1b (left) and 1e (left), a pairwise image similarity matrix (IxI) is precomputed for each image dataset with DreamSim. Each corresponding representation evaluation task directly queries this matrix to perform MEI comparisons.

To perform localized interpretability evaluations (Extended Data Figs. 1a (right), 1e (right)), one precomputed DreamSim similarity matrix is calculated per mask *M*, in addition to the image-wide similarity matrix. For example, the Chang et al. [39] dataset with *I* = 2100 images and *M* = 5 mask types (eyes, forehead, chin, outer face, inner face) yields 6 total 2100*X*2100 precomputed matrices and the best of 6 corresponding Odd-One-Out scores is saved per neuron. The values for the Majaj et al. [13] V4 dataset are *I* = 3200*, M* = 1 (mask type: image center).

### Image similarity metric: Wasserstein Distortion

Grounded in models of human vision, the Wasserstein Distortion [43] is a metric that can evaluate local image similarity without masking or cropping images by manipulating the parameter *σ* to simulate different receptive field sizes. Briefly, each point in an image is associated with a distribution of features around it. The weights associated with those features decrease as a function of distance from the point being evaluated. The rate of this weight decay is controlled by *σ*. For example, a small *σ* value leads to rapid weight fall-off, thereby defining the similarity between two images at that point more locally, as would a neuron with a small receptive field. While beneficial for localization of representation evaluations, it’s worth noting that default Wasserstein Distortion does not use human-aligned ensemble of models characteristic of Dream-Sim, but rather VGG-19 feature map activations to define the features (though see [43] for modifications).

Before measuring interpretability in Extended Data Fig. 1b (right), a pairwise image similarity matrix (IxI; on a subset of images) is precomputed from the Majaj et al. [13] images for three values of *σ* with the Wasserstein Distortion. The best Odd-One-Out value for each neuron is compared to a fixed Odd-One-Out value for a given (largest) receptive field size.

### Measuring Interpretability: Odd-One-Out Score

The Odd-One-Out task measures how distinguishable a given neuron or population’s *K* maximally exciting images (MEIs) are from the remaining images in the dataset. To compute this score for a single neuron (*A*), all images in the dataset are sorted in descending order according to their activation level for the neuron being evaluated. The top *K* images of this sorted list are the neuron’s MEIs, and their average pairwise similarity is computed to establish a task threshold:

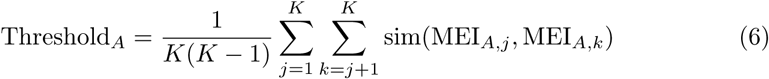

where sim(MEI*_A,j_,* MEI*_A,k_*) denotes the precomputed DreamSim or Wasserstein Distortion similarity between MEI pairs. Next, we calculate the average similarity of each of the remaining *D* images (where *D* = *I − K*) to the MEIs:

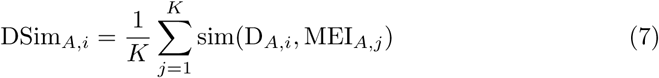

where sim(D*_A,i_,* MEI*_A,j_*) represents the precomputed DreamSim or Wasserstein Distortion similarity between D*_A,i_* and MEI*_A,j_*. One task point is assigned for each instance where DSim*_A,i_* < Threshold*_A_*, indicating the model identified the ‘odd-image-out’ among the MEIs + D*_A,i_*:

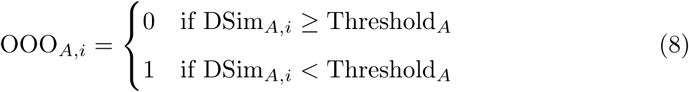

The final Odd-One-Out score for a neuron is the average across all *D* tasks. We repeat this procedure for all neurons and report the average. The same method applies to populations.

### Measuring Diversity: Cross Odd-One-Out Score

The Cross Odd-One-Out task measures the diversity of features represented across *N* neurons or *Z* populations by comparing the MEIs of all pairs of neurons or populations in a dataset. To obtain a Cross Odd-One-Out score for a pair of neurons (*A*, *B*), *K* MEIs from each unit are selected. The average pairwise similarity between each of the resulting 2*K* MEIs and 1) the MEIs of neuron A and 2) the MEIs of neuron B is computed, excluding the MEI being evaluated. The procedure to evaluate a single MEI from neuron *A* is as follows. First, similarity to other MEIs from neuron *A* is computed:

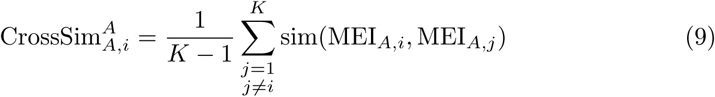

where sim(*·*) denotes the precomputed DreamSim similarity between MEIs. Next, similarity to MEIs from neuron *B* are computed:

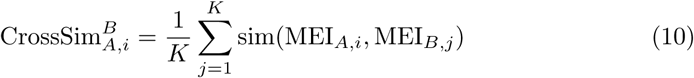

One point is scored for each instance that this MEI is more similar to the remaining MEIs of unit *A*, its unit of origin:

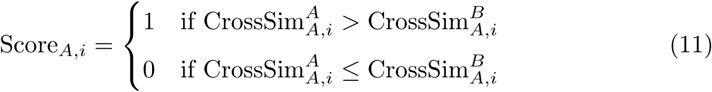

Repeating this process for the remaining MEIs of neuron *A* and neuron *B* and taking the average of all 2*K* tasks yields the final Cross Odd-One-Out score for neurons *A* and *B*. All other pairwise comparisons between neurons fill the remaining entries in the NxN Cross Odd-One-Out matrix. Note that this matrix is symmetric (e.g., the final score for the pair (*A*, *B*) equals the final score for the pair (*B*, *A*)). The same protocol extends to populations to generate a ZxZ Cross Odd-One-Out matrix.

### Quantifying visual feature yield

#### Subsampling and cluster expansion

To test how visual feature discovery scales with neural dataset size, biological neurons are randomly subsampled in intervals of 10 from 10 to 30 and artificial neurons are randomly subsampled in intervals of 10 from 10 to 100. Neuron selection is without replacement within, but not between, subsamples. To additionally test how visual feature discovery scales with the number of populations, four values of *Z* are chosen for each set of subsampled neurons *N*, given by *Z ∈ {N,* 2*N,* 4*N,* 8*N}*.

#### Greedy feature selection algorithm

A greedy algorithm reports the number of unique, interpretable visual features represented. First, *N* subsampled neurons or *Z* populations are sorted in descending order according to their Odd-One-Out scores. All representations with Odd-One-Out scores below 0.8 are discarded. Next, each interpretable neuron or population is appended to a list provided that it is unique, i.e., its Cross Odd-One-Out score with any unit already on the list does not fall below 0.8. The length of the resulting list is the number of interpretable, unique visual features represented across *N* neurons or *Z* populations. 0.8 was selected as the Odd-One-Out and Cross Odd-One-Out threshold based on visual confirmation that MEIs of greedy algorithm list members appear readily interpretable and diverse across datasets.

#### Label recovery

While MEIs corresponding to each interpretable, diverse feature from *N* neurons versus *Z* populations can be shown for biological neurons (Fig. 4h), the greedy feature selection algorithm returns a much larger feature set for artificial vision networks. Thus, label recovery is performed after greedy selection for the models. This method applies an additional layer by appending greedy algorithm outputs to a list only if 7 of 10 of their MEIs correspond to a single CIFAR-100 category and this category is not already represented in the list. The analysis is performed separately for coarse (20) and fine (100) CIFAR-100 labels. The list is cumulative, growing across subsampled neurons (10-100) or across corresponding populations at Z=8N (80-800).

#### Image dataset size and diversity

To quantify visual feature yield as a function of image dataset size, the number of sampled neurons is fixed at N=30 and the number of corresponding UTM codes is fixed at Z=240. 100 CIFAR-100 image categories are maintained, and the number of images per category is increased from 40-100 in steps of 20. To quantify visual feature yield as a function of image diversity, N=30 and Z=240, but 100 images per category are maintained, and the number of image categories increases from 40-100 in steps of 20. Across experiments, the total number of images at each step is identical. The greedy algorithm is applied to data at each step to determine the number of unique, interpretable features discovered.

### Ablation experiments

#### DINOv2

Targeted ablations of DINOv2 representations are used to test their causal contributions to downstream behavior. To define ‘neurons’ for ablation, *N* = 384 spatially pooled DINOv2 neural responses to CIFAR-100 images are collected then passed through 1) the greedy selection algorithm and 2) a purity filter requiring 7/10 MEIs to share a single category label. To identify populations for ablation, *Z* = 384 UTM clusters were fit to per-patch DINOv2 responses to CIFAR-100 images. UTM clusters are similarly passed through the greedy selection algorithm and category purity filter. Next, a five-way classification task is automatically defined for each discovered feature by identifying the four additional object categories in its CIFAR-100 Super-class. DINOv2 classification performance before and after feature ablation is reported. Classification uses a weighted *k*-nearest-neighbor vote (cosine similarity, temperature-weighted) over L2-normalized, unmodified DINOv2 embeddings of held-out images, following standard DINOv2 evaluation protocol. UTM feature ablations project raw neural activations of DINOv2 onto the UTM cluster center and subtract the projection. Neuron ablations remove the contribution of the ablated neuron from raw DINOv2 neural activations. In addition to precise causal manipulations, this procedure enables observation of image regions most impacted, offering post-hoc explanations for ablation performance.

#### IT Cortex

To extend this causal framework to biological neurons, macaque IT neurons [35] and population responses are ablated using the same gain-control logic. First, a correspondence between IT neurons (or populations) and DINOv2 is established to enable the same post-hoc patch-based activation visualizations as the previous section. After identifying diverse, interpretable, purity-filtered DINOv2 UTM clusters (as above, but computed on responses to the IT stimulus set), linear regression predicts DINOv2 UTM responses from single neurons to define IT neurons for ablation (without replacement via Hungarian algorithm). Multivariate linear regression predicts DINOv2 UTM responses from the entire IT population to define populations for ablation. Only neurons or populations with *R*^2^ *≥* 0.60 are retained for the ablation experiment, which follows the same structure as the DINOv2 ablation experiment except a linear classifier trained to predict image categories directly from IT population activity is used in absence of animal behavior.

### Feature synergy

Null neurons are generated from the original data by taking the most interpretable IT cortex neuron (according to the Odd-One-Out task), duplicating it 85 times to match the original number of neurons for subject 1 of the Vinken et al. [35] dataset, and adding independent Gaussian noise equal to the standard deviation of the original neurons. This yields a null population whose baseline interpretability is comparable to that of the real population. The change in interpretability and diversity of this population going from neurons to UTM codes is subsequently compared to the same change on the original neurons.

### Statistics

Kruskal-Wallis tests with Dunn’s post-hoc pairwise comparisons (Bonferroni-corrected) measure mean rank differences in interpretability (Fig. 2) and diversity (Fig. 3) scores. All other statistical comparisons are made using paired Student’s t-tests.

## Acknowledgments

This work was performed with assistance from the US National Institutes of Health Grant S10OD028632-01.

## Extended Data

**Fig. S1:**
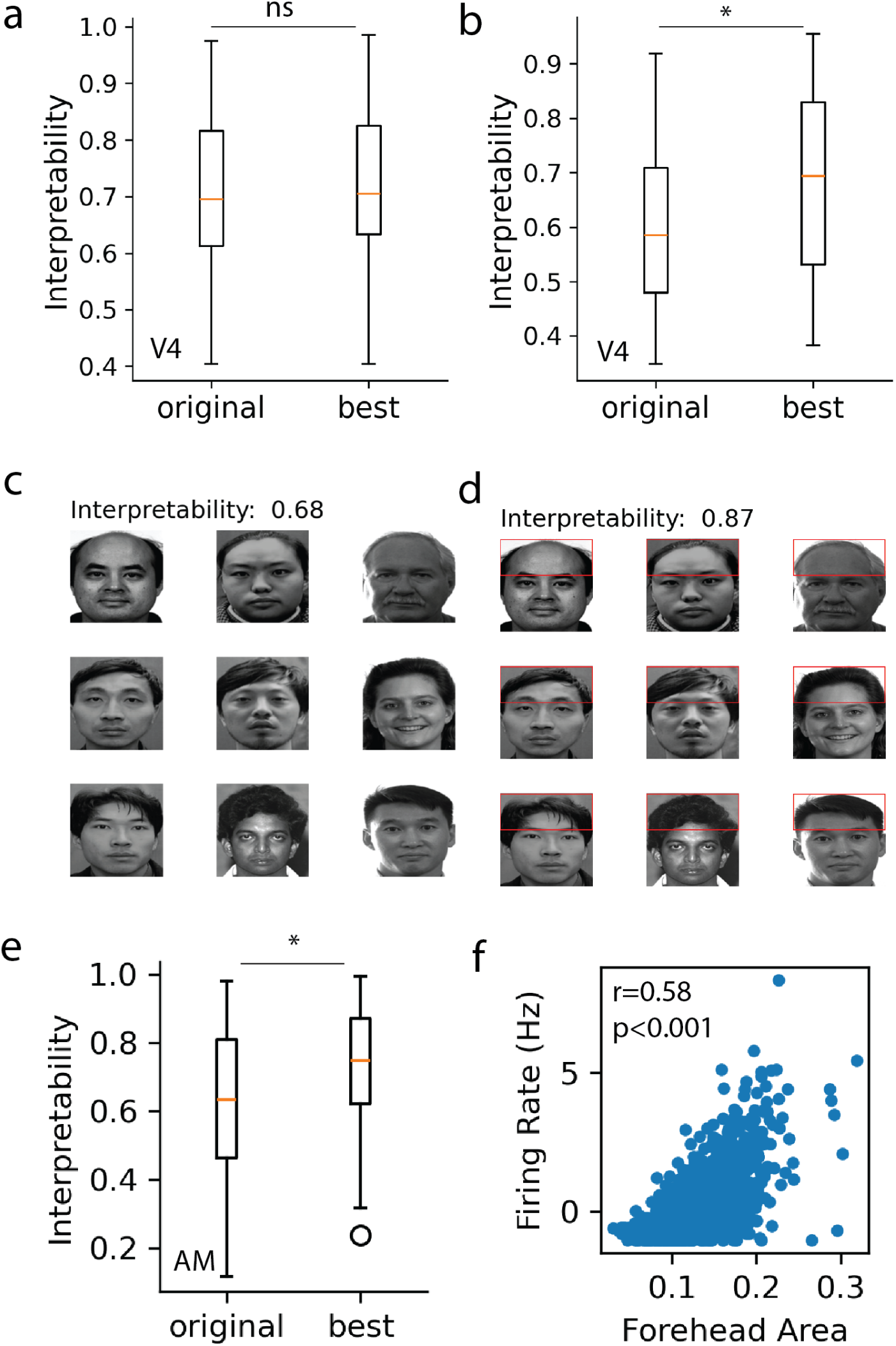
Local interpretability measurements. **a**, Original Odd-One-Out score for V4 neurons compared to best local Odd-One-Out score with DreamSim. **b,** Original Odd-One-Out score for V4 neurons with Wasserstein Distortion compared to best local Odd-One-Out score with Wasserstein Distortion. **c,** Odd-One-Out score and MEIs for example AM unit with image-wide Dreamsim. Images from Chang et al. [39]. **d,** Odd-One-Out score and MEIs for example AM unit with DreamSim measurements localized to forehead. Images from Chang et al. [39]. **e,** Original Odd-One-Out score for all AM neurons compared to best Odd-One-Out score from local DreamSim measurements **f,** Correlation between example AM unit and forehead area, measured with MediaPipe [58]. ns = Not Significant, *∗* = *P <* 0.05

**Fig. S2:**
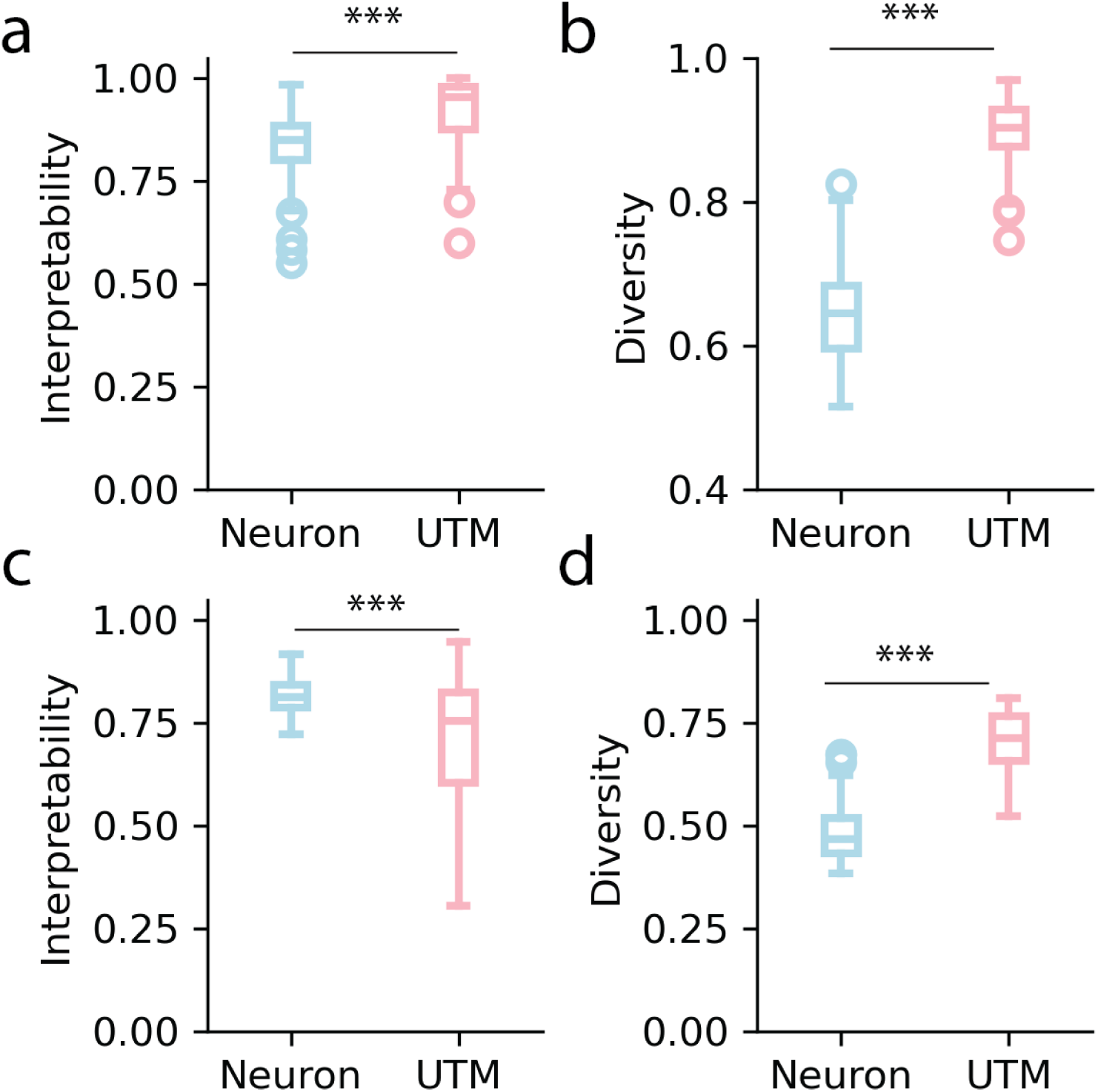
Population synergy. **a**, Odd-One-Out scores for real neurons versus corresponding UTM codes. **b,**Cross Odd-One-Out scores for real neurons versus corresponding UTM codes. **c,** Odd-One-Out scores for null neurons versus corresponding UTM codes. **d,** Cross Odd-One-Out scores for null neurons versus corresponding UTM codes. ns = Not Significant, *∗* = *P <* 0.05, *∗ ∗ ∗* = *P <* 0.001

**Fig. S3:**
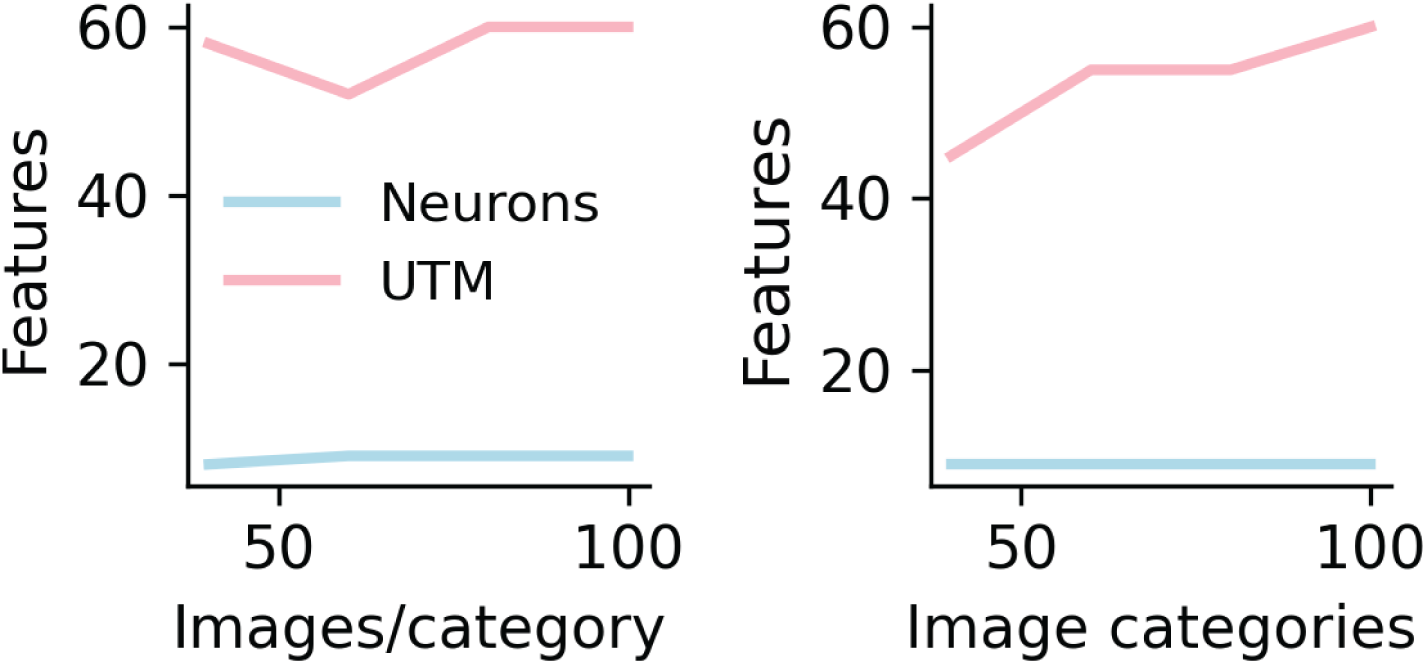
Image diversity. Number of discovered features as a function of images per category (left) and images categories (right).

**Fig. S4:**
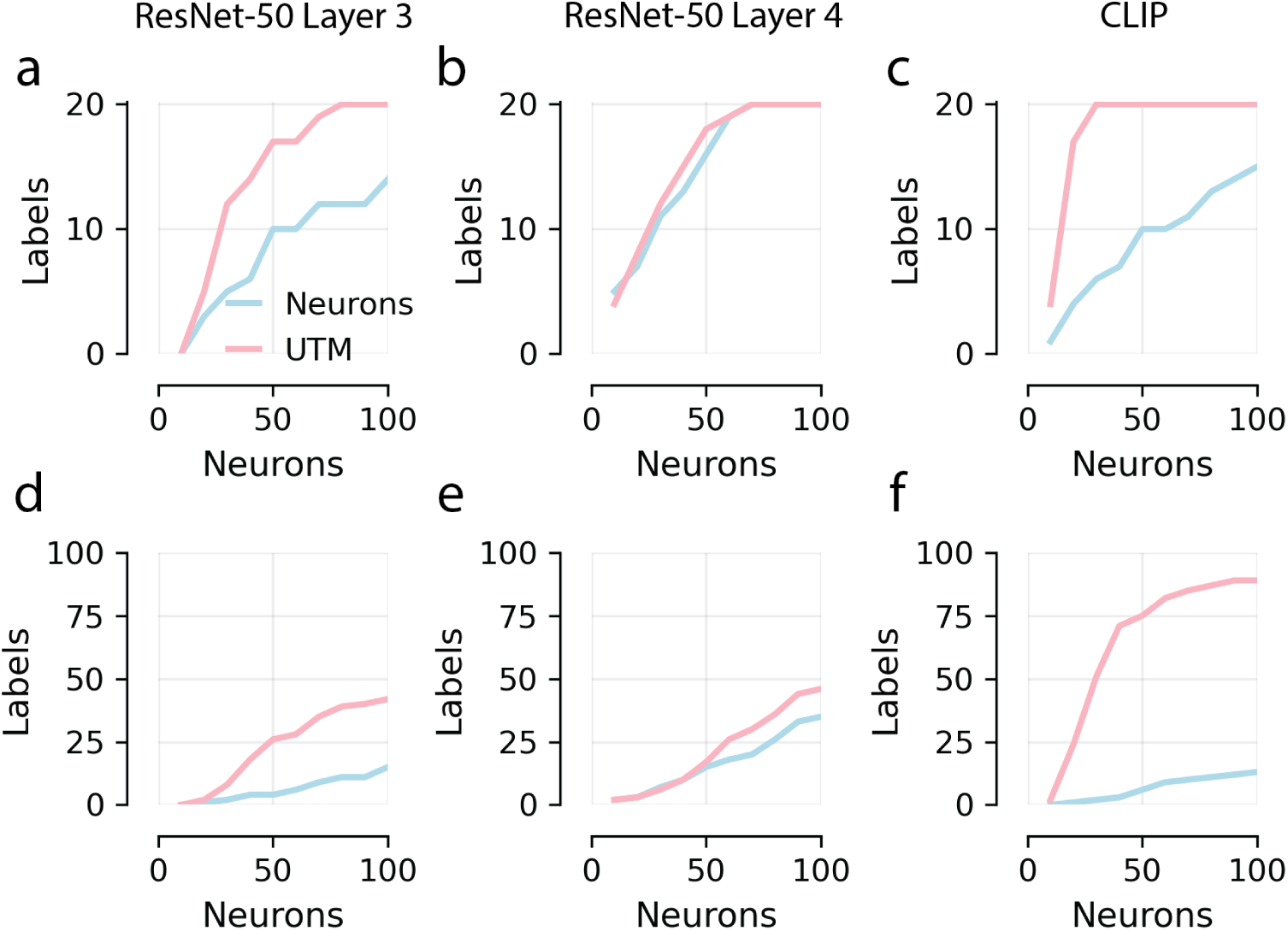
Label recovery. **a-c** Number of neurons (blue) versus UTM codes (pink) representing a coarse CIFAR-100 label as a function of number of sample neurons for ResNet-50 layer 3 (**a**), ResNet-50 layer 4 (**b**), and CLIP (**c**). **d-f** Number of neurons (blue) versus UTM codes (pink) representing a fine CIFAR-100 label as a function of number of sample neurons for ResNet-50 layer 3 (**d**), ResNet-50 layer 4 (**e**), and CLIP (**f**).

---

**Table 1:** Odd-One-Out Statistics: Kruskal-Wallis.

| ID | N | df | H | p |
| --- | --- | --- | --- | --- |
| Vinken2023_IT_3 | 30 | 3 | 14.0 | 0.00291 |
| Vinken2023_IT_5 | 44 | 3 | 37.27 | < 0.001 |
| MajajHong2015_IT_0 | 58 | 3 | 75.52 | < 0.001 |
| MajajHong2015_V4_0 | 70 | 3 | 74.46 | < 0.001 |
| Vinken2023_IT_2 | 85 | 3 | 135.94 | < 0.001 |
| MajajHong2015_IT_1 | 110 | 3 | 154.49 | < 0.001 |
| Vinken2023_IT_0 | 267 | 3 | 555.98 | < 0.001 |
| CLIP_Cifar100 | 512 | 3 | 1377.54 | < 0.001 |
| ResNet50_layer3_Cifar100 | 1024 | 3 | 2660.69 | < 0.001 |
| ResNet50_layer4_Cifar100 | 2048 | 3 | 5891.19 | < 0.001 |

**Table 2:** Odd-One-Out: Dunn’s Test Post-Hoc Comparisons.

| ID | Comparison | N | Mean Diff | Cohen's d_z | p-adj |
| --- | --- | --- | --- | --- | --- |
| Vinken2023_IT_3 | Neuron_PCA | 30 | 0.09 | 0.49 | 0.247 |
| Vinken2023_IT_3 | Neuron_SAE | 30 | -0.03 | -0.21 | 1 |
| Vinken2023_IT_3 | PCA_SAE | 30 | -0.12 | -0.85 | 0.00416 |
| Vinken2023_IT_3 | Neuron_UTM | 30 | -0.02 | -0.13 | 1 |
| Vinken2023_IT_3 | SAE_UTM | 30 | 0.01 | 0.05 | 1 |
| Vinken2023_IT_3 | PCA_UTM | 30 | -0.11 | -0.58 | 0.0134 |
| Vinken2023_IT_5 | Neuron_UTM | 44 | -0.02 | -0.15 | 0.86 |
| Vinken2023_IT_5 | PCA_SAE | 44 | -0.09 | -0.53 | < 0.001 |
| Vinken2023_IT_5 | Neuron_PCA | 44 | 0.13 | 0.84 | < 0.001 |
| Vinken2023_IT_5 | Neuron_SAE | 44 | 0.04 | 0.26 | 1 |
| Vinken2023_IT_5 | SAE_UTM | 44 | -0.05 | -0.33 | 0.259 |
| Vinken2023_IT_5 | PCA_UTM | 44 | -0.14 | -0.81 | < 0.001 |
| MajaJHong2015_IT_0 | Neuron_SAE | 58 | -0.03 | -0.16 | 1 |
| MajaJHong2015_IT_0 | Neuron_PCA | 58 | 0.18 | 0.91 | < 0.001 |
| MajaJHong2015_IT_0 | PCA_SAE | 58 | -0.21 | -0.96 | < 0.001 |
| MajaJHong2015_IT_0 | Neuron_UTM | 58 | -0.1 | -0.55 | < 0.001 |
| MajaJHong2015_IT_0 | SAE_UTM | 58 | -0.07 | -0.39 | 0.0218 |
| MajaJHong2015_IT_0 | PCA_UTM | 58 | -0.28 | -1.22 | < 0.001 |
| MajaJHong2015_V4_0 | Neuron_PCA | 70 | 0.13 | 0.64 | < 0.001 |
| MajaJHong2015_V4_0 | Neuron_SAE | 70 | -0.02 | -0.11 | 1 |
| MajaJHong2015_V4_0 | SAE_UTM | 70 | -0.07 | -0.33 | 0.0551 |
| MajaJHong2015_V4_0 | PCA_UTM | 70 | -0.22 | -1.18 | < 0.001 |
| MajaJHong2015_V4_0 | PCA_SAE | 70 | -0.15 | -0.85 | < 0.001 |
| MajaJHong2015_V4_0 | Neuron_UTM | 70 | -0.09 | -0.51 | 0.00196 |
| Vinken2023_IT_2 | Neuron_SAE | 85 | -0.02 | -0.19 | 0.135 |
| Vinken2023_IT_2 | Neuron_PCA | 85 | 0.17 | 0.82 | < 0.001 |
| Vinken2023_IT_2 | PCA_UTM | 85 | -0.25 | -1.27 | < 0.001 |
| Vinken2023_IT_2 | SAE_UTM | 85 | -0.06 | -0.44 | 0.00263 |
| Vinken2023_IT_2 | Neuron_UTM | 85 | -0.08 | -0.72 | < 0.001 |
| Vinken2023_IT_2 | PCA_SAE | 85 | -0.2 | -0.93 | < 0.001 |
| MajaJHong2015_IT_1 | SAE_UTM | 110 | -0.06 | -0.38 | 0.0881 |
| MajaJHong2015_IT_1 | PCA_UTM | 110 | -0.31 | -1.58 | < 0.001 |
| MajaJHong2015_IT_1 | Neuron_PCA | 110 | 0.21 | 0.87 | < 0.001 |
| MajaJHong2015_IT_1 | Neuron_SAE | 110 | -0.04 | -0.23 | 0.359 |
| MajaJHong2015_IT_1 | Neuron_UTM | 110 | -0.1 | -0.65 | < 0.001 |
| MajaJHong2015_IT_1 | PCA_SAE | 110 | -0.25 | -1.12 | < 0.001 |
| Vinken2023_IT_0 | SAE_UTM | 267 | -0.16 | -0.92 | < 0.001 |
| Vinken2023_IT_0 | PCA_UTM | 267 | -0.39 | -2.05 | < 0.001 |
| Vinken2023_IT_0 | Neuron_PCA | 267 | 0.24 | 1.07 | < 0.001 |
| Vinken2023_IT_0 | Neuron_SAE | 267 | 0.01 | 0.05 | 1 |
| Vinken2023_IT_0 | Neuron_UTM | 267 | -0.15 | -0.93 | < 0.001 |
| Vinken2023_IT_0 | PCA_SAE | 267 | -0.23 | -0.93 | < 0.001 |
| CLIP_Cifar100 | Neuron_SAE | 512 | -0.12 | -0.76 | < 0.001 |
| CLIP_Cifar100 | Neuron_PCA | 512 | 0.24 | 1.28 | < 0.001 |
| CLIP_Cifar100 | PCA_UTM | 512 | -0.42 | -2.93 | < 0.001 |
| CLIP_Cifar100 | SAE_UTM | 512 | -0.06 | -0.61 | < 0.001 |
| CLIP_Cifar100 | Neuron_UTM | 512 | -0.18 | -1.52 | < 0.001 |
| CLIP_Cifar100 | PCA_SAE | 512 | -0.35 | -2.09 | < 0.001 |
| ResNet50_layer3_Cifar100 | Neuron_PCA | 1024 | 0.45 | 2.9 | < 0.001 |
| ResNet50_layer3_Cifar100 | Neuron_SAE | 1024 | 0.0 | 0.02 | 0.379 |
| ResNet50_layer3_Cifar100 | PCA_UTM | 1024 | -0.54 | -4.16 | < 0.001 |
| ResNet50_layer3_Cifar100 | SAE_UTM | 1024 | -0.09 | -0.84 | < 0.001 |
| ResNet50_layer3_Cifar100 | PCA_SAE | 1024 | -0.45 | -2.69 | < 0.001 |
| ResNet50_layer3_Cifar100 | Neuron_UTM | 1024 | -0.09 | -0.91 | < 0.001 |
| ResNet50_layer4_Cifar100 | Neuron_UTM | 2048 | -0.08 | -0.9 | < 0.001 |
| ResNet50_layer4_Cifar100 | PCA_SAE | 2048 | -0.38 | -2.73 | < 0.001 |
| ResNet50_layer4_Cifar100 | Neuron_SAE | 2048 | 0.02 | 0.17 | < 0.001 |
| ResNet50_layer4_Cifar100 | Neuron_PCA | 2048 | 0.4 | 3.08 | < 0.001 |
| ResNet50_layer4_Cifar100 | PCA_UTM | 2048 | -0.48 | -4.82 | < 0.001 |
| ResNet50_layer4_Cifar100 | SAE_UTM | 2048 | -0.1 | -1.03 | < 0.001 |

**Table 3:** Cross Odd-One-Out Statistics: Kruskal-Wallis.

| ID | N | df | H | p |
| --- | --- | --- | --- | --- |
| Vinken2023_IT_3 | 30 | 3 | 77.58 | < 0.001 |
| Vinken2023_IT_5 | 44 | 3 | 126.91 | < 0.001 |
| MajaHong2015_IT_0 | 58 | 3 | 149.92 | < 0.001 |
| MajaHong2015_V4_0 | 70 | 3 | 152.0 | < 0.001 |
| Vinken2023_IT_2 | 85 | 3 | 265.11 | < 0.001 |
| MajaHong2015_IT_1 | 110 | 3 | 251.22 | < 0.001 |
| Vinken2023_IT_0 | 267 | 3 | 838.03 | < 0.001 |
| CLIP_Cifar100 | 512 | 3 | 1745.34 | < 0.001 |
| ResNet50_layer3_Cifar100 | 1024 | 3 | 3194.62 | < 0.001 |
| ResNet50_layer4_Cifar100 | 2048 | 3 | 6872.3 | < 0.001 |

**Table 4:** Cross Odd-One-Out: Dunn’s Test Post-Hoc Comparisons.

| ID | Comparison | N | Mean Diff | Cohen's d_z | p-adj |
| --- | --- | --- | --- | --- | --- |
| Vinken2023_IT_3 | Neuron_PCA | 30 | 0.02 | 0.14 | 1 |
| Vinken2023_IT_3 | Neuron_SAE | 30 | -0.15 | -1.86 | < 0.001 |
| Vinken2023_IT_3 | PCA_SAE | 30 | -0.17 | -1.77 | < 0.001 |
| Vinken2023_IT_3 | Neuron_UTM | 30 | -0.19 | -2.02 | < 0.001 |
| Vinken2023_IT_3 | SAE_UTM | 30 | -0.04 | -0.38 | 1 |
| Vinken2023_IT_3 | PCA_UTM | 30 | -0.2 | -2.31 | < 0.001 |
| Vinken2023_IT_5 | Neuron_SAE | 44 | -0.14 | -1.36 | < 0.001 |
| Vinken2023_IT_5 | Neuron_PCA | 44 | 0.05 | 0.46 | 0.565 |
| Vinken2023_IT_5 | PCA_SAE | 44 | -0.19 | -2.13 | < 0.001 |
| Vinken2023_IT_5 | Neuron_UTM | 44 | -0.21 | -2.43 | < 0.001 |
| Vinken2023_IT_5 | SAE_UTM | 44 | -0.07 | -1.22 | 0.00504 |
| Vinken2023_IT_5 | PCA_UTM | 44 | -0.26 | -3.09 | < 0.001 |
| MajaHong2015_IT_0 | Neuron_SAE | 58 | -0.05 | -0.69 | 0.0035 |
| MajaHong2015_IT_0 | Neuron_PCA | 58 | 0.11 | 1.46 | < 0.001 |
| MajaHong2015_IT_0 | PCA_SAE | 58 | -0.15 | -1.96 | < 0.001 |
| MajaHong2015_IT_0 | Neuron_UTM | 58 | -0.1 | -1.24 | < 0.001 |
| MajaHong2015_IT_0 | SAE_UTM | 58 | -0.05 | -0.76 | 0.0047 |
| MajaHong2015_IT_0 | PCA_UTM | 58 | -0.2 | -2.24 | < 0.001 |
| MajaHong2015_V4_0 | PCA_UTM | 70 | -0.17 | -1.73 | < 0.001 |
| MajaHong2015_V4_0 | SAE_UTM | 70 | -0.05 | -0.54 | 0.0266 |
| MajaHong2015_V4_0 | Neuron_UTM | 70 | -0.08 | -0.94 | < 0.001 |
| MajaHong2015_V4_0 | PCA_SAE | 70 | -0.13 | -1.55 | < 0.001 |
| MajaHong2015_V4_0 | Neuron_SAE | 70 | -0.04 | -0.45 | 0.0311 |
| MajaHong2015_V4_0 | Neuron_PCA | 70 | 0.09 | 1.05 | < 0.001 |
| Vinken2023_IT_2 | SAE_UTM | 85 | -0.09 | -1.26 | < 0.001 |
| Vinken2023_IT_2 | PCA_UTM | 85 | -0.3 | -3.19 | < 0.001 |
| Vinken2023_IT_2 | PCA_SAE | 85 | -0.21 | -2.27 | < 0.001 |
| Vinken2023_IT_2 | Neuron_UTM | 85 | -0.25 | -3.2 | < 0.001 |
| Vinken2023_IT_2 | Neuron_PCA | 85 | 0.05 | 0.46 | 0.156 |
| Vinken2023_IT_2 | Neuron_SAE | 85 | -0.17 | -1.85 | < 0.001 |
| MajaHong2015_IT_1 | Neuron_SAE | 110 | -0.03 | -0.39 | 0.0159 |
| MajaHong2015_IT_1 | Neuron_PCA | 110 | 0.18 | 1.82 | < 0.001 |
| MajaHong2015_IT_1 | PCA_UTM | 110 | -0.24 | -2.38 | < 0.001 |
| MajaHong2015_IT_1 | SAE_UTM | 110 | -0.03 | -0.51 | 0.0113 |
| MajaHong2015_IT_1 | Neuron_UTM | 110 | -0.06 | -0.87 | < 0.001 |
| MajaHong2015_IT_1 | PCA_SAE | 110 | -0.2 | -2.18 | < 0.001 |
| Vinken2023_IT_0 | Neuron_PCA | 267 | 0.21 | 2.27 | < 0.001 |
| Vinken2023_IT_0 | Neuron_SAE | 267 | 0.01 | 0.15 | 0.986 |
| Vinken2023_IT_0 | SAE_UTM | 267 | -0.18 | -2.02 | < 0.001 |
| Vinken2023_IT_0 | PCA_UTM | 267 | -0.38 | -4.46 | < 0.001 |
| Vinken2023_IT_0 | Neuron_UTM | 267 | -0.17 | -2.34 | < 0.001 |
| Vinken2023_IT_0 | PCA_SAE | 267 | -0.2 | -1.74 | < 0.001 |
| CLIP_Cifar100 | Neuron_SAE | 512 | -0.09 | -1.82 | < 0.001 |
| CLIP_Cifar100 | Neuron_PCA | 512 | 0.18 | 2.48 | < 0.001 |
| CLIP_Cifar100 | PCA_UTM | 512 | -0.31 | -4.51 | < 0.001 |
| CLIP_Cifar100 | SAE_UTM | 512 | -0.04 | -0.98 | < 0.001 |
| CLIP_Cifar100 | Neuron_UTM | 512 | -0.13 | -3.58 | < 0.001 |
| CLIP_Cifar100 | PCA_SAE | 512 | -0.27 | -3.67 | < 0.001 |
| ResNet50_layer3_Cifar100 | Neuron_PCA | 1024 | 0.31 | 4.95 | < 0.001 |
| ResNet50_layer3_Cifar100 | Neuron_SAE | 1024 | -0.0 | -0.02 | 1 |
| ResNet50_layer3_Cifar100 | Neuron_UTM | 1024 | -0.06 | -1.7 | < 0.001 |
| ResNet50_layer3_Cifar100 | PCA_SAE | 1024 | -0.32 | -4.45 | < 0.001 |
| ResNet50_layer3_Cifar100 | SAE_UTM | 1024 | -0.06 | -1.39 | < 0.001 |
| ResNet50_layer3_Cifar100 | PCA_UTM | 1024 | -0.38 | -5.89 | < 0.001 |
| ResNet50_layer4_Cifar100 | PCA_SAE | 2048 | -0.31 | -4.81 | < 0.001 |
| ResNet50_layer4_Cifar100 | Neuron_UTM | 2048 | -0.07 | -1.94 | < 0.001 |
| ResNet50_layer4_Cifar100 | SAE_UTM | 2048 | -0.09 | -2.31 | < 0.001 |
| ResNet50_layer4_Cifar100 | PCA_UTM | 2048 | -0.4 | -7.5 | < 0.001 |
| ResNet50_layer4_Cifar100 | Neuron_SAE | 2048 | 0.02 | 0.39 | < 0.001 |
| ResNet50_layer4_Cifar100 | Neuron_PCA | 2048 | 0.33 | 5.34 | < 0.001 |

